# Correlating STED and synchrotron XRF nano-imaging unveils the co-segregation of metals and cytoskeleton proteins in dendrites

**DOI:** 10.1101/810754

**Authors:** Florelle Domart, Peter Cloetens, Stéphane Roudeau, Asuncion Carmona, Emeline Verdier, Daniel Choquet, Richard Ortega

**Author notes:** D.C. and R.O. share seniority.

## Abstract

Zinc and copper are involved in neuronal differentiation and synaptic plasticity but the molecular mechanisms behind these processes are still elusive due in part to the difficulty of imaging trace metals together with proteins at the synaptic level. We correlate stimulated emission depletion (STED) microscopy of proteins and synchrotron X-ray fluorescence (SXRF) imaging of trace metals, both performed with 40 nm spatial resolution, on primary rat hippocampal neurons. We achieve a detection limit for zinc of 14 zeptogram (10^-21^ g) per pixel. We reveal the co-localization at the nanoscale of zinc and tubulin in dendrites with a molecular ratio of about one zinc atom per tubulin-αβ dimer. We observe the co-segregation of copper and F-actin within the nano-architecture of dendritic protrusions. In addition, zinc chelation causes a decrease in the expression of cytoskeleton proteins in dendrites and spines. Overall, these results indicate new functions for zinc and copper in the modulation of the cytoskeleton morphology in dendrites, a mechanism associated to neuronal plasticity and memory formation.

---

The neurobiology of copper and zinc is a matter of intense investigation since they have been recently associated to neuronal signaling and differentiation processes (Chang, 2015; Vergnano et al., 2014; Barr et al., 2017; D’Ambrosi et al., 2015; Xiao et al., 2018; Hatori et al., 2016). Zinc ions are stored in pre-synaptic vesicles in a subset of glutamatergic neurons. They are released in the synaptic cleft during neurotransmission. Zinc ions are effectors of synaptic plasticity by modulating the activity of neurotransmitter receptors such as NMDA (N-methyl-d-aspartate) and AMPA (α-amino-3-hydroxy-5-methyl-4-isoxazolepropionic acid) receptors (NMDARs and AMPARs) (Vergnano et al., 2014; Barr et al., 2017). The role of copper in neural functions is less well described than that of zinc, but copper ions are also known to be released in the synaptic cleft where they can modulate the activity of neurotransmitter receptors (D’Ambrosi et al., 2015). Copper is involved in broad neuronal functions such as the regulation of spontaneous neuronal activity or of rest–activity cycles (Chang, 2015; Xiao et al., 2018), and more generally in neuronal differentiation (Hatori et al., 2016). In addition to these functions of the labile fraction of metals on neuronal signaling and differentiation, we have recently indicated that copper and zinc could be involved in the structure of dendrites and dendritic spines (Perrin et al., 2017). We have shown on cultured hippocampal neurons the localization of zinc all along dendritic shafts and that of copper in the neck of dendritic spines, suggesting their interaction with cytoskeleton proteins. Understanding the functions of copper and zinc in the cytoskeleton structure would require however to correlate metal localization with respect to relevant cytoskeleton proteins at the sub-dendritic level. To this end, we have developed an original method for correlative imaging of metals and proteins at 40 nm spatial resolution described in this paper.

Synchrotron X-Ray Fluorescence (SXRF) imaging using hard X-ray beams, typically above 10 keV energy, is a powerful technique to investigate the cellular localization of metals since it allows the mapping of element distributions in single cells with high analytical sensitivity (Pushie et al., 2014). Using Kirkpatrick–Baez (KB) focusing mirrors, SXRF has reached a record spatial resolution of 13 nm on ID16A beamline at the European Synchrotron Radiation Facility (ESRF), while maintaining a high photon flux as required for detecting trace elements (Da Silva et al., 2017). We have previously reported a correlative microscopy approach consisting in labeling organelles or proteins with specific fluorophores for live-cell imaging prior to SXRF imaging (Roudeau et al., 2014; Carmona et al., 2019). This correlative approach is limited by the spatial resolution of optical fluorescence microscopy, above 200 nm, which is larger than the spatial resolution achieved today with nano-SXRF and insufficient to resolve synaptic sub-structures. To overcome this limitation we present a method to correlate nano-SXRF with STED (STimulated Emission Depletion microscopy) performed both at 40 nm resolution. With the combination of these two high resolution imaging techniques we observe trace metals co-localization with cytoskeleton proteins at the synaptic level in rat hippocampal neurons. Furthermore, we explored the effect of zinc deficiency induced by TPEN, an intracellular zinc chelator, on the expression of cytoskeleton proteins in dendrites and dendritic spines.

## Results

### Super resolution live-cell STED microscopy and nano-SXRF imaging combination

We designed a specific protocol consisting in live-cell STED microscopy on silicon nitride (SN) substrates followed by cryogenic processing of the cells before nano-SXRF imaging as described in (fig. 1). Primary rat hippocampal neurons were cultured *in vitro* during 15 days (DIV15) on sterile SN membranes placed above an astrocyte feeder layer as adapted from Kaech & Banker (Kaech & Banker, 2006; Perrin et al., 2015). SN membranes were designed with an orientation frame to facilitate the reproducible positioning of the samples during the correlative imaging procedure (fig. 1a). SN membranes are biocompatible, trace metal free, 500 nm thin, transparent, flat and rigid supports developed for X-ray microscopy. To perform STED microscopy we stained the DIV15 primary rat hippocampal neurons with either SiR-actin, SiR-tubulin, or with SiR-tubulin and SiR700-actin together. These two far-red SiR-based fluorophores have been developed for live-cell super resolution microscopy and are based respectively on the actin ligand jasplakinolide and the tubulin ligand docetaxel (Lukinavičius et al., 2014; Lukinavičius et al., 2016). Cytoskeletal SiR-based fluorescent dyes have several features that were important for the success of this correlative imaging approach. On the one hand, they are markers suitable for STED imaging that do not require overexpression of the proteins. On the other hand, when these molecules bind to proteins of the cytoskeleton they block the cell structure preventing movement of the regions of interest in the time span between living cells STED imaging and cryofixation. Confocal and STED live-cell images were obtained using a commercial Leica DMI6000 TCS SP8 X with a 93x objective at immersion in glycerol and numerical aperture of 1.3 (fig. 1b). STED images were recorded at 40 nm spatial resolution. For each region of interest (roi) the (x,y) coordinates were registered in the confocal/STED microscope setup. Then the SN membranes were plunge-frozen at −165°C and freeze-dried at −90°C under secondary vacuum before analysis on ID16A beamline at ESRF (fig. 1b). Using the coordinates of 3 reference points on the SN membrane (3 corners of the membrane relatively to the orientation frame), the new coordinates (x’,y’) of the roi selected during STED microscopy were calculated for the ID16A microscope setup (fig. 1a). Nano-SXRF and synchrotron X-ray phase contrast imaging (PCI) were performed with an X-ray beam size of 40 nm at 17 KeV energy (fig. 1b). Nano-SXRF, PCI, and STED microscopy images can be merged to perform multimodal correlative microscopy (fig. 1c) of chemical elements, electron density, and fluorescently labeled proteins all obtained at similar spatial resolution (<40 nm). Post-acquisition alignment of the multimodal images was performed using intra-specimen localization structures such as characteristic filament crosses, or dendritic protrusions. The result is accurate despite the freeze drying process that can marginally alter the cell structure (fig. 1c). The adequacy of the sample preparation method and of the post-acquisition alignment of the images for the correlative purpose was checked by comparing live cell STED with STXM (Scanning Transmission X-ray Microscopy) of freeze dried cells obtained with 25 nm spatial resolution at HERMES beamline, SOLEIL synchrotron. STXM enabled to visualize the dendritic morphology showing the exact superposition of dendrites after cryofixation and freeze drying with SiR-tubulin fluorescence previously observed by STED on living cells (supplementary fig. S1). The limit of detection (LOD) of nano-SXRF analysis was calculated for the detected elements according to IUPAC (International Union of Pure and Applied Chemistry) guideline resulting for zinc in in a 0.009 ng.mm^-2^ LOD, corresponding to 14 zeptogram of zinc (about 130 atoms) per pixel of 40 nm x 40 nm size (Table 1).

**Figure 1.**
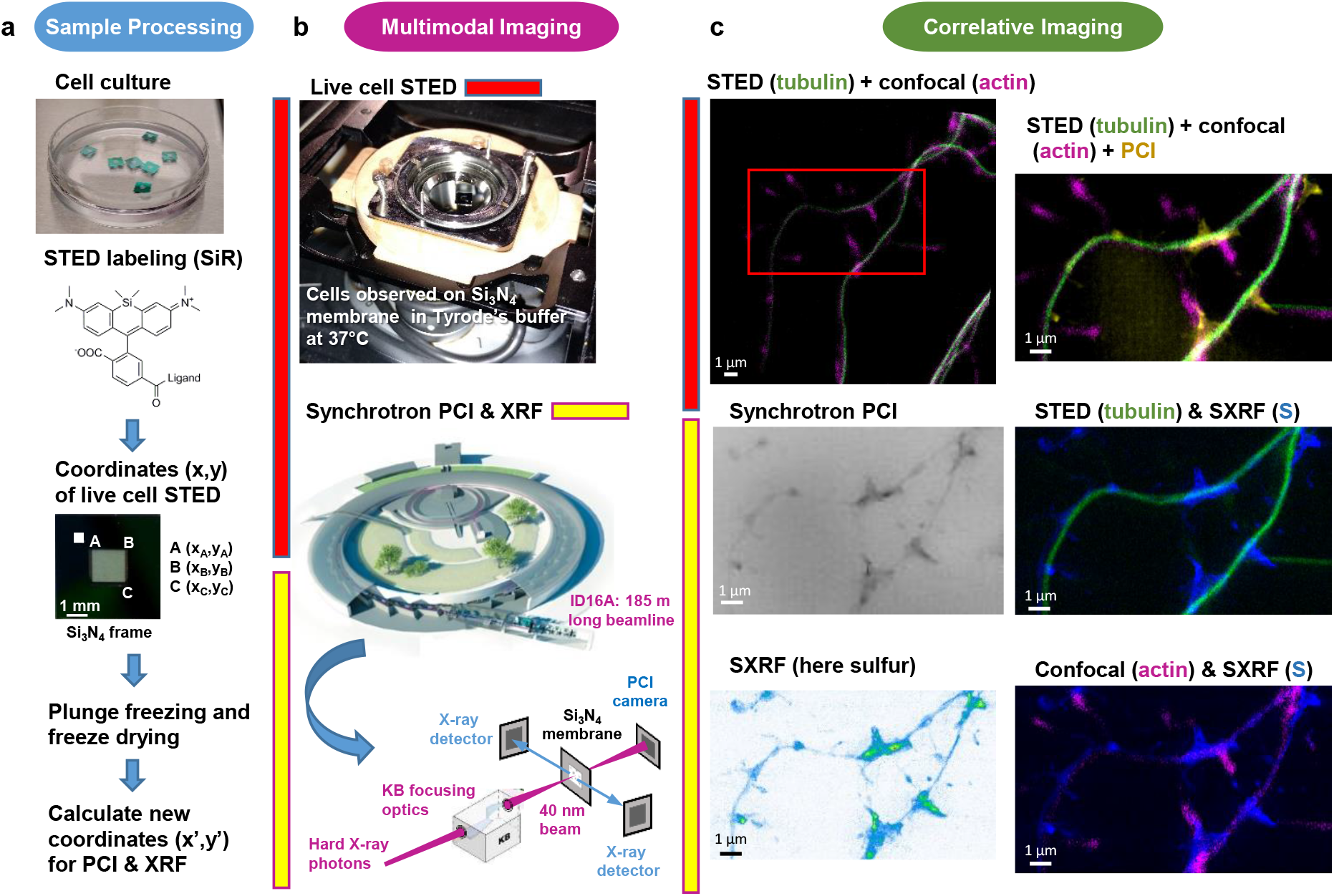
Workflow for correlative STED and synchrotron multimodal nano-imaging. **a**, Sample processing. Primary neurons are cultured on silicon nitride membranes and labelled with fluorescent probes designed for STED microscopy such as SiR-tubulin or SiR700-actin. STED microscopy is performed on living cells and orthonormal coordinates (x,y) of regions of interest are recorded relatively to the position of three membrane corners A, B, C identified thanks to an orientation frame. Immediately after STED microscopy cells are plunge-frozen and freeze-dried. New coordinates (x’,y’) of the regions of interest are calculated to perform XRF and PCI imaging on the synchrotron microscope. **b**, Multi-modal imaging. Live-cell STED and confocal microscopy are performed with a Leica DMI6000 TCS SP8 X microscope equipped with a thermalized chamber. Synchrotron XRF and PCI are carried out on freeze-dried samples at ESRF on beamline ID16A. The KB optics are 185 m away from the X-ray source enabling to focus hard X-rays at 40 nm beam size. **c**, Correlative imaging. Overlay images of STED, confocal, synchrotron PCI and XRF are produced on areas of few tens of μm large with a spatial resolution of 40 nm for STED, 30 nm for PCI and 40 nm for SXRF. Several elemental maps (here sulfur) can be super-imposed with protein distributions (i.e. actin or tubulin) in dendrites and spines at 40 nm spatial resolution.

**Table 1.** Nano-SXRF limit of detection (LOD). LOD calculated according to IUPAC (International Union of Pure and Applied Chemistry) guideline as derived from 12 blank measurements for phosphorus, sulfur, potassium and zinc, expressed in ng.mm^-2^, and expressed in g (and number of atoms) within pixels of 40 nm x 40 nm size.

|  | Phosphorus | Sulfur | Potassium | Zinc |
| --- | --- | --- | --- | --- |
| **LOD** | 0.513 ng.mm<sup>-2</sup> | 0.138 ng.mm<sup>-2</sup> | 0.152 ng.mm<sup>-2</sup> | 0.009 ng.mm<sup>-2</sup> |
| **LOD in 40 nm** | 8.2 10<sup>-19</sup> g | 2.2 10<sup>-19</sup> g | 2.4 10<sup>-19</sup> g | 1.4 10<sup>-20</sup> g |
| **x 40 nm pixel** | (16,000 atoms) | (4,200 atoms) | (3,800 atoms) | (130 atoms) |

### Zinc is highly concentrated in dendritic spines

The highest zinc content is found in F-actin-rich dendritic spines where zinc is highly correlated with sulfur (Person’s correlation coefficient > 0.6) (fig. 2–4 and S2–S4). Zinc distribution however is not fully superimposed with F-actin localization suggesting that both entities are not directly in interaction in dendritic protrusions. Zinc and sulfur hot spots are found within narrow regions in the dendritic spines characterized by the highest electron densities as shown by synchrotron PCI maps (fig. 3 and S4). Although in lower concentration than zinc, copper was also detected in F-actin rich dendritic spines showing a distinct distribution pattern than for Zn (fig. 3 and S3). Copper is mainly located at the basis of the dendritic protrusions (fig. 3h) where it is co-localized with the highest fluorescence signal of F-actin (fig. 3j). The co-localization of copper and F-actin in dendritic protrusions is only partial since copper may not be detected in regions of weaker SiR-actin fluorescence. In rare cases iron was also detected, as 200 nm x 100 nm structures of locally dense iron spots, in the dendrite, contiguous to the dendritic spines (fig. S4K).

**Figure 2.**
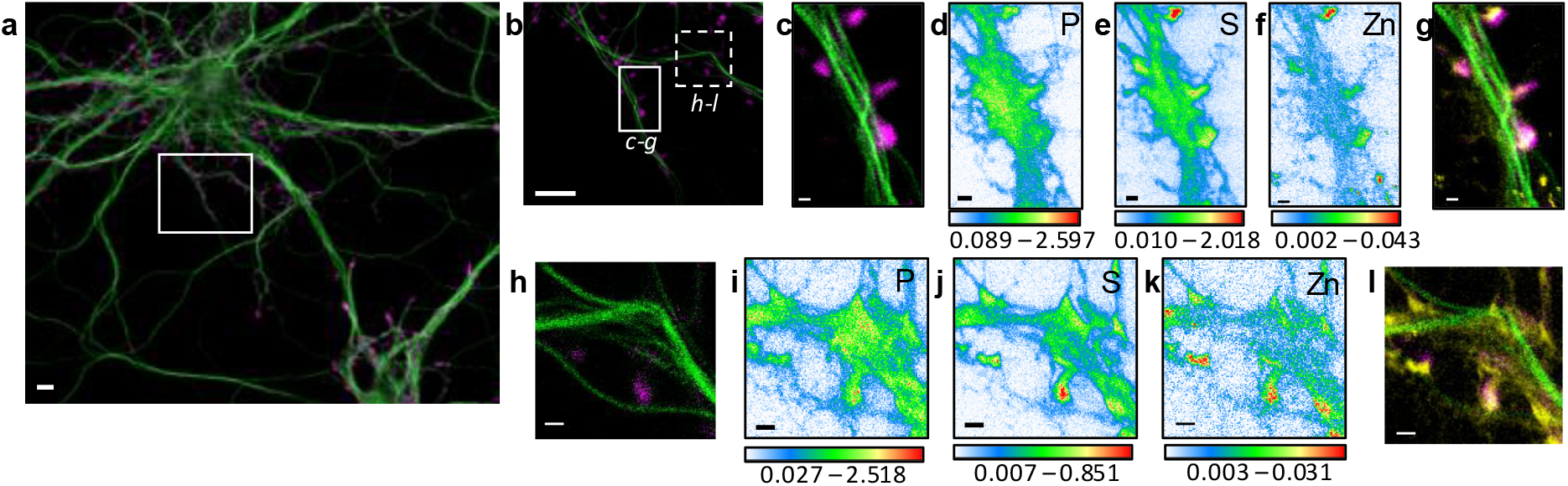
Correlative STED nano-SXRF element imaging in dendritic spines. **a**, Confocal image from a DIV15 primary rat hippocampal neuron stained with SiR-tubulin (green) and SiR700-actin (magenta). **b**, STED image of SiR-tubulin (green) and confocal SiR700-actin (magenta) from the framed region shown in (**a**). **c**, STED image of SiR-tubulin (green) and confocal SiR700-actin (magenta) for the region mapped by SXRF shown in (**b**) in plain line. **d**, Phosphorus SXRF map. **e**, Sulfur SXRF map. **f**, Zinc SXRF map. **g**, Merged images of zinc SXRF map (yellow), STED SiR-tubulin (green) and confocal SiR700-actin (magenta). **h**, STED image of SiR-tubulin (green) and confocal SiR700-actin (magenta) image from the framed region shown in (**b**) in dotted line. **i,**) Phosphorous SXRF map. **j**, Sulfur SXRF map. **k**, Zinc SXRF map. **l**, Merged images of zinc SXRF map (yellow), STED SiR-tubulin (green), and confocal SiR700-actin (magenta). Scale bars: 500 nm, except (**a**) and (**b**) 5 μm. Color scale bars: min-max values in ng.mm^-2^.

**Figure 3.**
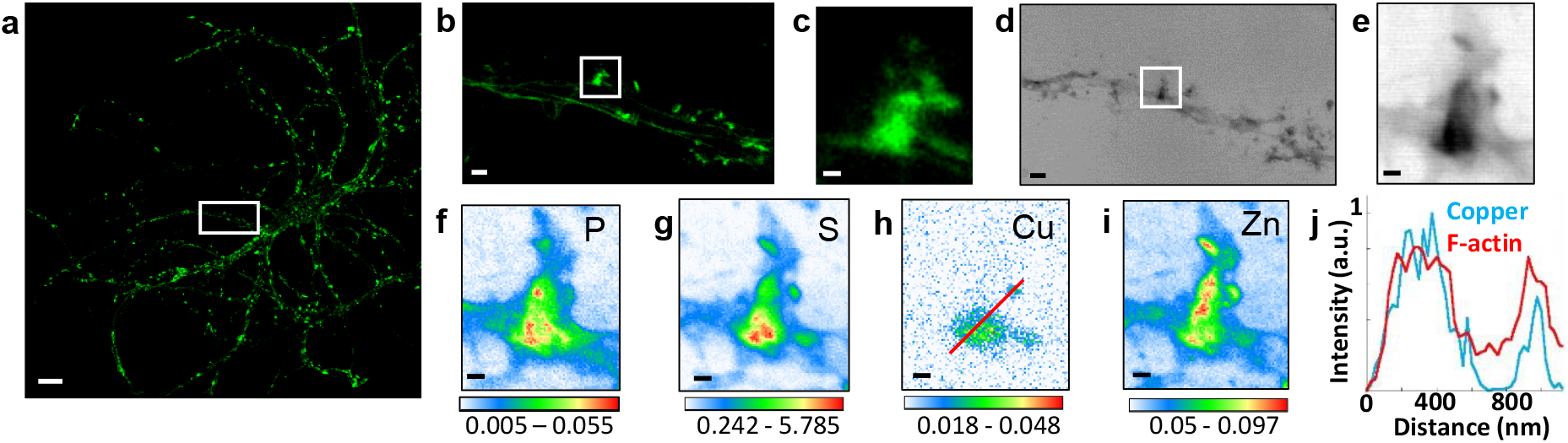
Correlative STED microscopy of SiR-actin with synchrotron PCI and XRF in spines. **a**, Confocal imaging of a primary rat hippocampal neuron labeled with SiR-actin showing Factin-rich protrusions along dendrites. **b**, STED imaging of SiR-actin of the dendrite region framed in white in A. **c**, Zoom on the STED SiR-actin of a F-actin-rich protrusion framed in (**b**). **d**, Synchrotron radiation X-ray PCI of the dendrite region framed in white in (**a**). **e**, Zoom on the PCI region of a F-actin-rich protrusion framed in (**d**). **f-i**, SXRF element maps (P, S, Cu, Zn) from the region of interest framed in (**b**) and (**d**). **j**, Line scans for F-actin (red) and copper (blue) normalized distributions along the red line plotted in (**h**). Scale bar: 200 nm, except for (**a**) 10 μm, (**b**) and (**d**) 1 μm. Color scales: min-max values in ng.mm^-2^.

### Zinc and microtubules are co-localized

The most striking result of this correlative microscopy approach is the observation of zinc and tubulin co-localization in thin dendritic processes as illustrated in fig. 4–5 and S3–S5. In fig. 5, a roi showing parallel microtubule filaments observed by STED microscopy was selected (fig. 5a). These parallel tubulin filaments could not be resolved by confocal microscopy (fig. 5b), but only by STED microscopy (fig. 5c). Similarly to STED microscopy, nano-SXRF performed on the same roi was able to separate the element distributions of the two thin dendritic processes (fig. 5d-e). Plot profiles of zinc and tubulin relative signal intensities across the two tubulin filaments shows the co-localization of their distributions (fig. 5f). The very good superimposition of S and Zn distributions along tubulin imaged by STED is systematically observed (fig. 4, 5k and Fig. S3–S5). For thinner dendrites, zinc is superimposed with the dendritic structure, while for thicker branches zinc is located in sub-dendritic regions (supplementary fig. S5). The quantitative analysis of nano-SXRF data for 21 regions showing Zn and tubulin co-localization indicates that the atomic ratio S/Zn is of 44 ± 11 (fig. 5L and Table S1). Considering that the amino acid sequence of the rat tubulin-α/β dimer contains 49 sulfur atoms per dimer, from methionine and cysteine residues, the S/Zn ratio in microtubules corresponds to a theoretical tubulin-α/β dimer over Zn ratio of 0.9 ± 0.2 (fig. 5l).

**Figure 4.**
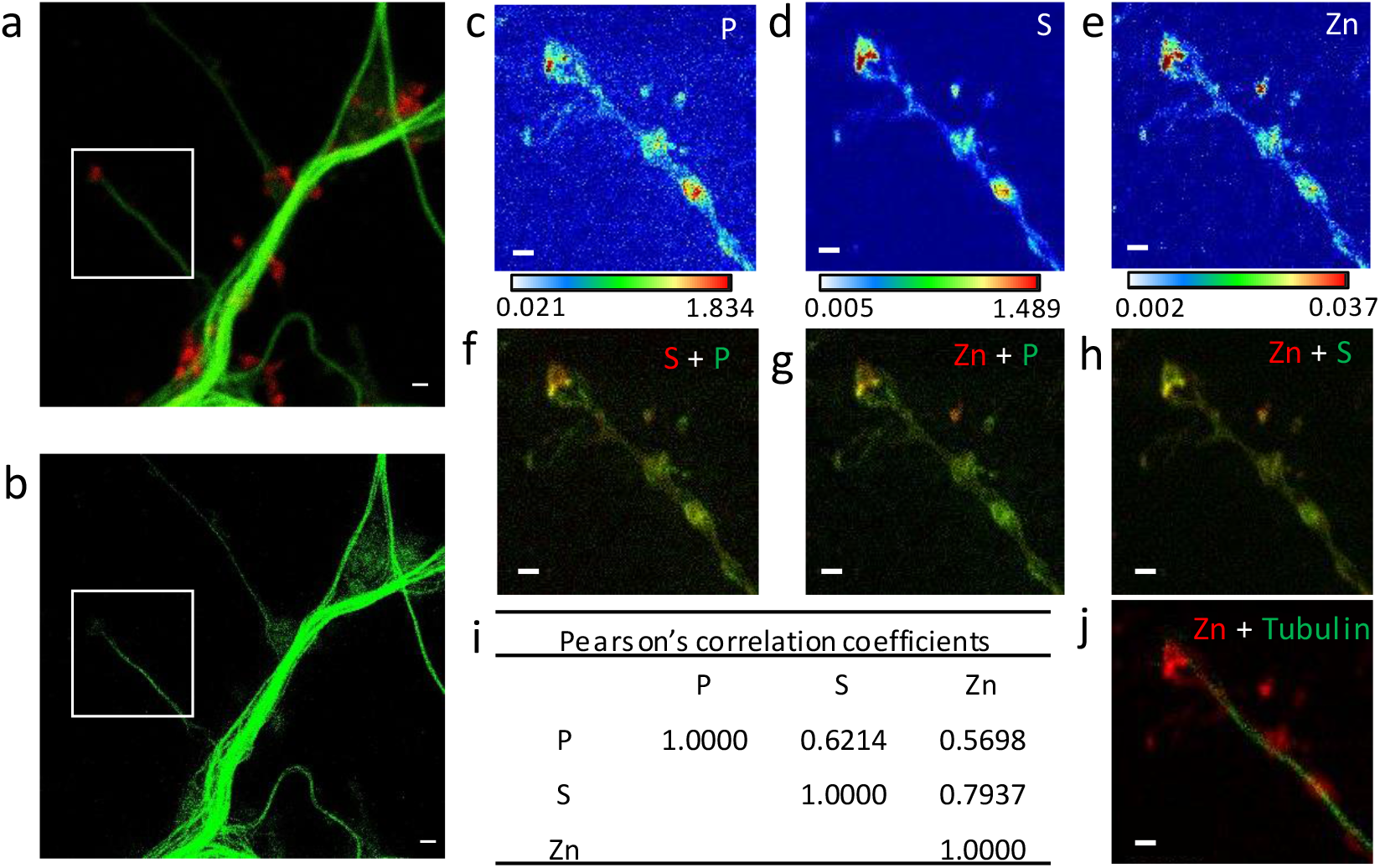
Correlative imaging in dendrites and spines. **a**, Confocal image of SiR-tubulin (green) and SiR700-actin (red). **b**, STED image of SiR-tubulin (green) for the same zone as in (**a**). **c-e**, SXRF element maps (P, S, Zn) from the region of interest framed in (**a**) and (**b**). **f**, Overlay image of phosphorus (green) and sufur (red). **g**, Overlay image of phosphorus (green) and zinc (red). **h**, Overlay image of sulfur (green) and zinc (red). **i**, Pearson’s correlation coefficients for the elements (P, S, Zn) in the roi. **j**, Overlay image of STED SiR-tubulin (green) and zinc (red). Scale bars: 500 nm, except for (**a**) and (**b**) 1 μm. Color scales min-max values in ng.mm^-2^.

**Figure 5.**
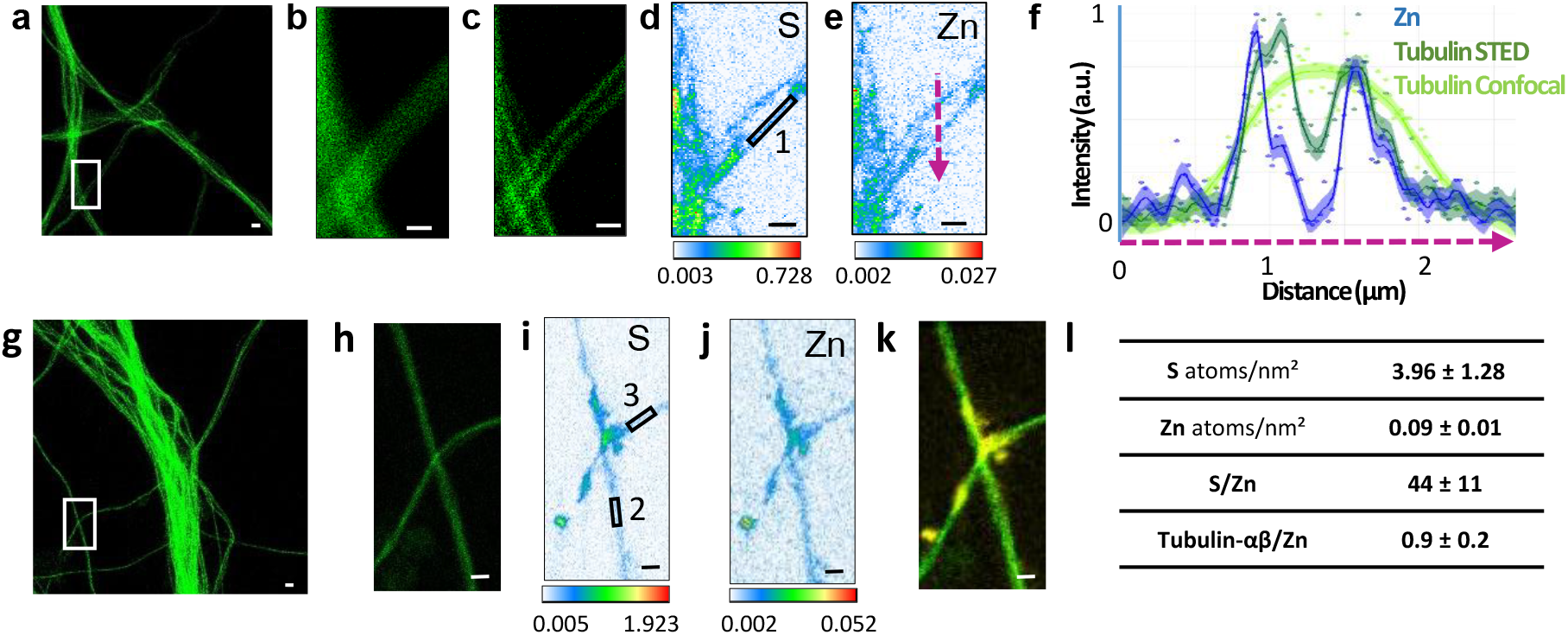
Correlative STED and nano-SXRF imaging in dendrites. **a**, STED image of SiR-tubulin (green). **b**, Confocal SiR-tubulin image of the region framed in (**a**). **c**, STED SiR-tubulin image of the region framed in (**a**). **d**, Sulfur SXRF map. **e**, Zinc SXRF map. **f**, Intensity plot profiles of zinc (blue), STED SiR-tubulin (dark green), and confocal SiR-tubulin (light green) along a line scan crossing two thin dendrites (magenta dotted line in (**e**)). **g**, STED image of SiR-tubulin (green) in dendrites. **h**, STED SiR-tubulin of the region framed in (**g**). **i**, Sulfur SXRF map. **j**, Zinc SXRF map. **k**, Overlay image of STED SiR-tubulin (green) and zinc distribution (yellow). **l**, Quantitative data analysis of the number of sulfur and zinc atoms.nm^-2^ for 21 regions of interest centered on thin microtubules as illustrated for roi 1 to 3 framed in D and I (mean ± SD; n=21; see also Table S1). Scale bars: 500 nm, except for (**a**) and (**g**) 1 μm.

### Zinc depletion disturbs the dendritic cytoskeleton

Primary rat hippocampal neurons cultured on glass coverslips were exposed to TPEN (N,N,N’,N’-tetrakis(2-pyridylmethyl)ethylenediamine), a zinc intracellular chelator. At DIV 15, mature neurons were exposed to a subcytotoxic concentration of 5 μM TPEN during 24h, or simultaneously to 5 μM TPEN and 10 μM ZnCl_2_, or to 0.1% ethanol (v/v) for the control group. Subcytotoxic concentration of TPEN during 24h exposure time was selected to study the effect of mild zinc depletion conditions on neuronal differentiation, avoiding cytoskeleton alteration that would result from direct cytotoxic effects at higher TPEN concentrations. Quantitative fluorescence microscopy of β-tubulin and F-actin expression was assessed in dendritic branches.

Representative examples of F-actin and β-tubulin labelling in dendrites for each of the three experimental groups are shown in fig. 6a-c. Fluorescence intensities of F-actin and β-tubulin were quantified in dendritic branches from more than 60 different neurons and were normalized with respect to the median value of the control group (fig. 6d-e). The experiment was repeated on three biological replicates (Fig. S6). Zinc chelation with 5 μM TPEN results in a statistically significant decrease of β-tubulin and F-actin fluorescence compared to the control group (fig. 6d-e and S6). The decrease in β-tubulin and F-actin expression following TPEN exposure is reversed by the addition of Zn strongly suggesting that the effect of TPEN is due to Zn chelation and not to unspecific interactions (fig. 6d-e and S6).

**Figure 6.**
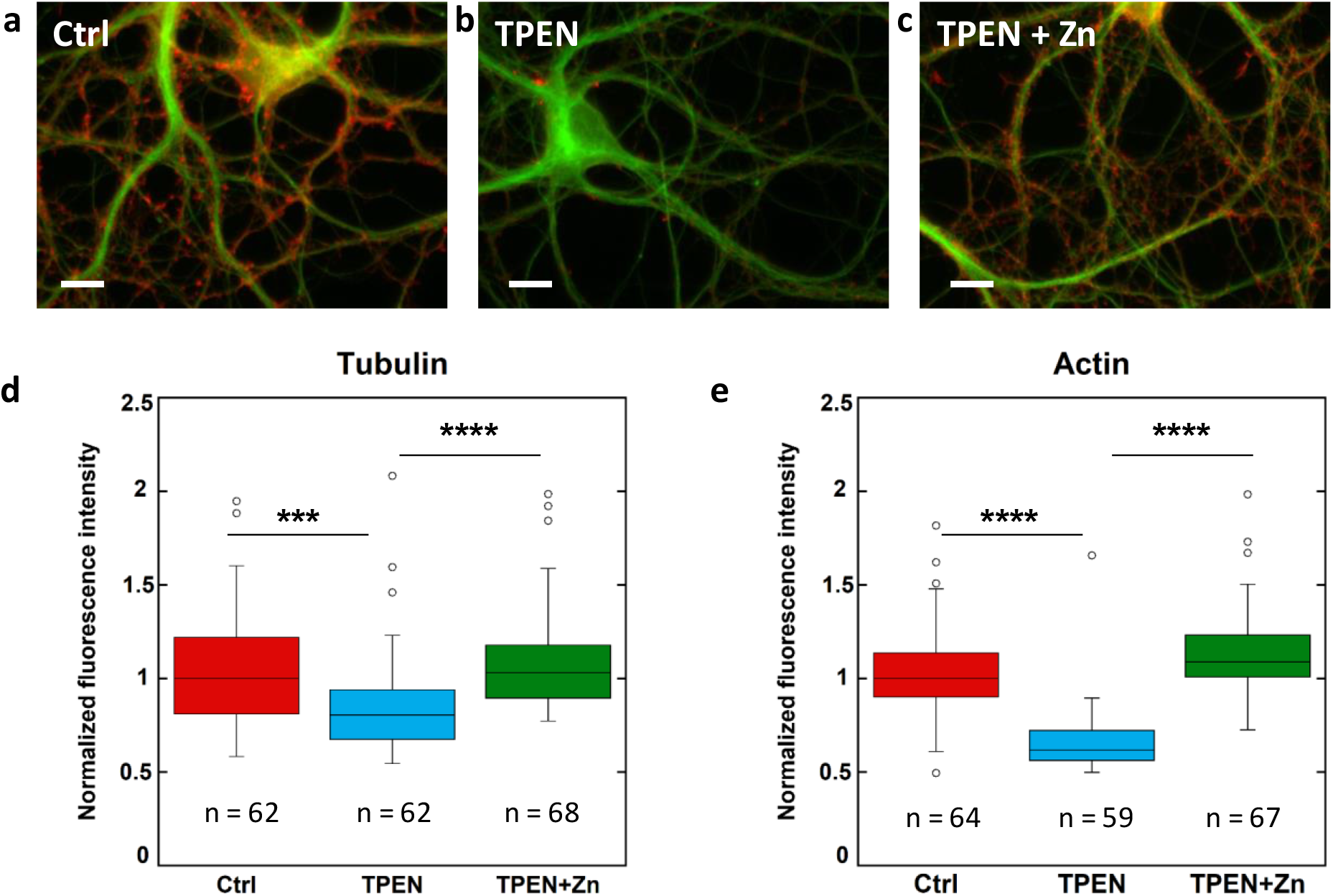
β-tubulin and F-actin fluorescence in control, TPEN, or TPEN and Zn treated neurons. **a**, Representative fluorescence microscopy images of F-actin (red) and β-tubulin (green) in dendrites from control neurons. Scale bar = 10 μm. **b**, TPEN treated neurons (5 μM, 24h). **c**, TPEN (5 μM, 24h) and Zn (10 μM, 24h) treated neurons. **d**, Comparison of β-tubulin normalized fluorescence intensities. **e**, Comparison of F-actin normalized fluorescence intensities. Significant Kruskal-Wallis test (p<0.01) was followed by Dunn’s test with p-values adjusted for pairwise comparisons: ***adj.p-value <0.001, ****adj.p-value <0.0001 (detailed in Table S2).

## Discussion

Zinc and copper are essential metals for neuronal functions (Chang, 2015; Vergnano et al., 2014; Barr et al., 2017; D’Ambrosi et al., 2015; Xiao et al., 2018; Hatori et al., 2016). Zinc is present in different regions of the brain, mainly the hippocampus and the amygdala (Frederickson et al., 2000). Zinc is largely complexed to proteins and is required for the functioning of hundreds of proteins. Only 10% of the total zinc is in a labile state, often referred as zinc ions (Zn^2+^), a more easily exchangeable and therefore mobile element pool (Takeda, 2001). In a subset of glutamatergic neurons, this labile zinc is highly concentrated at the pre-synaptic level where zinc is stored in synaptic vesicles (Vergnano et al., 2014; Barr et al., 2017). As for other neurotransmitters, zinc is released in the synaptic cleft where it inhibits neuronal transmission mediated by AMPARs or NMDARs (Vergnano et al., 2014; Barr et al., 2017; Kalappa et al., 2015). Zinc dyshomeostasis is broadly associated to brain diseases such as age-related cognitive decline, depression, or Alzheimer’s disease (Portbury et al., 2017). Copper is another essential metal for various cellular functions such as respiration, defense against free radicals and neurotransmitter synthesis (Chang, 2015; D’Ambrosi et al., 2015; Xiao et al., 2018; Hatori et al., 2016). Contrary to zinc ions, copper is not localized within synaptic vesicles but copper ions can be released into the synaptic cleft by means of copper transporter proteins (D’Ambrosi et al., 2015). Copper can play a biphasic role on neural transmission mediated by AMPARs, either by blocking or increasing the neurotransmission depending on its concentration and time of exposure (Peters et al., 2011). As for zinc, a disturbed copper homeostasis is in found in many neurological diseases (D’Ambrosi et al., 2015). Copper deficiency is observed in brain regions targeted for neurodegeneration such as the substantia nigra in Parkinson’s disease (Davies et al., 2014), or the hippocampus in Alzheimer’s disease (Xu et al., 2017).

Due to their low concentration in cells, the investigation of zinc and copper functions requires sensitive analytical methods and, in particular, the identification of the proteins interacting with those metals remains a difficult analytical challenge. In this context, we combined two high-resolution chemical imaging methods, STED super resolution photonic fluorescence microscopy (Hell & Wichmann, 1994; Nägerl et al., 2008) and synchrotron XRF nano-imaging (Pushie et al., 2014) to gather information on protein and element distributions at the nanometer scale in cells. STED microscopy has already been successfully combined to transmission electron microscopy to correlate protein localization with cell ultrastructure (Watanabe et al., 2011), or to atomic force microscopy to investigate protein aggregation at high spatial resolution (Cosentino et al., 2019). Recently, STED has been performed together with synchrotron scanning diffraction microscopy to inform about diffraction patterns in cells (Bernhardt et al., 2018). These correlative approaches however cannot be transposed to the imaging of metals in cells since they require steps of chemical fixation known to disrupt the metal-binding equilibrium in cells (Roudeau et al., 2014; Perrin et al., 2015). Moreover, glass coverslips used for STED microscopy usually contain significant amounts of zinc and of some other trace metals. We designed an original protocol to perform SXRF and STED correlative microscopy that fulfils specific requirements in terms of substrate for cell culture and protocols for sample preparation to avoid element contamination, loss or redistribution. Our protocol combines high resolution imaging (40 nm) and high elemental sensitivity (zeptogram level). We focused our investigations on metal interactions with two cytoskeleton proteins, Factin and tubulin based on our previous SXRF nano-imaging results showing the localization of zinc in dendritic shafts and of copper in the spines of hippocampal neurons (Perrin et al., 2017).

Now, we observed the co-localization of zinc and tubulin in dendritic shafts at the nanometric level, including within the thinner dendrites. Moreover, we determined a sulfur to zinc molecular ratio of 44 ± 11, in regions of zinc and tubulin co-localization. Based on the known amino acid composition of neuronal rat tubulin-αβ dimer, with 49 sulfur atoms per dimer originating from 29 methionine and 20 cysteine amino acids, zinc co-localization with microtubules results in a stoichiometry of 0.9 ± 0.2 zinc atom per tubulin-αβ dimer. This result is in excellent agreement with data obtained by the Scatchard method on purified microtubule preparations from pig brains showing that zinc could bind tubulin with a molecular ratio of 0.86 ± 0.18 zinc atom per tubulin-αβ dimer (Hesketh, 1983). The Scatchard method consists in measuring the affinity of tubulin for zinc ions by adding known quantities of zinc to the purified protein. Our result is also in agreement with X-ray crystallography data of the tubulin-α/β dimer predicting the presence of one putative zinc ion per dimer, located in the tubulin-α subunit of zinc-induced tubulin sheets (Löwe et al., 2001). Our data provided by the correlation of STED and nano-SXRF imaging were obtained directly on primary cultured neurons in physiological conditions, without zinc addition, strongly supporting the biochemical hypothesis of a structural requirement for zinc in tubulin network assembly. Although our imaging approach cannot prove a direct interaction between zinc and tubulin, a molecular ratio of one zinc atom per tubulin-αβ dimer strongly suggests such a direct interaction.

There are other biochemical evidence indicating the interaction of tubulin and zinc. Immobilized metal affinity chromatography (IMAC) combined to mass spectrometry proteomic analysis has been applied to study the copper and zinc proteome of HepG2 human liver cancer cells (She et al., 2003). Nineteen copper-binding proteins and 19 zinc-binding proteins were identified. Interestingly, α-tubulin and β-tubulin are present among the 19 zinc binding-proteins identified by IMAC. Furthermore, in a proteomic analysis of HeLa cells lysates, based on the detection of proteins containing zinc-binding cysteines, both α-tubulin and β-tubulin were identified as putative zinc-binding proteins (Pace & Weerapana, 2014). These results may also apply to hippocampal neurons known to express α-tubulin and β-tubulin. Molecular modeling have identified six main binding sites of zinc to tubulin, suggesting that zinc would stabilize the structure of microtubules by enhancing electrostatic interactions (Craddock et al., 2012). Taken together, these previous biochemical and biophysical data (Hesketh, 1983; Craddock et al., 2012), and our results obtained in physiological conditions on hippocampal neurons provide a body of evidence suggesting a structural role for zinc in microtubules assembly.

The structural role of zinc on tubulin assembly could explain why this element is so fundamental for neuronal differentiation (Perrin et al., 2017; Chowanadisai et al., 2013), and why zinc dyshomeostasis is broadly involved in neuronal disorders. A reduction in microtubules size is observed in Alzheimer’s disease (Cash et al., 2003). It has been hypothesized that zinc dyshomeostasis could alter tubulin dynamics in Alzheimer’s disease (Craddock et al., 2013). In our study, this hypothesis is supported by the reduction of the neuronal cytoskeleton induced by zinc depletion using the intracellular zinc-chelator TPEN. We found that TPEN exposure resulted in decreased tubulin expression and that this effect was reversed by Zn. The decrease in tubulin expression can be explained by a limited neuronal differentiation resulting in a reduced density of tubulin filaments. Since F-actin expression in dendrites is an event that follows the development of tubulin network, the reduction in Factin dendritic protrusions is a consequence of the limited neuronal differentiation due to Zn chelation. These zinc chelation results are in agreement with our previous observation that zinc supplementation in the culture medium induces the increase of β-tubulin and F-actin expression in mature primary rat hippocampal neurons (Perrin et al., 2017). There is further evidence that zinc deficiency reduces tubulin expression in neurons. In neuronal cells cultured with a Zn depleted medium, as well as in brains of an animal model of Zn deficiency, a low rate of tubulin polymerization was observed (Mackenzie et al., 2011). In cultured human neuronal precursor NT2 cells, zinc deficiency decreased retinoic acid induced differentiation resulting in a large depletion of beta-tubulin(III) (Gower-Winter et al., 2013). More generally zinc deficiency impairs neuronal differentiation through a variety of mechanisms including processes such as DNA replication, transcriptional control, mRNA translation, apoptosis and microtubule stability (Pfaender et al., 2016). Overall these data suggest that zinc depletion leads to a decrease in tubulin expression through direct binding effects and indirect effects on protein regulation.

We also observe a concomitant decrease of F-actin expression that could be explained as the consequence of tubulin decrease since microtubules are involved in the regulation of dendritic spines through actin remodeling (Jaworski et al., 2009). The highest zinc content is found in dendritic spines, mainly located at the head of the spines, and zinc distribution was correlated with the higher cellular density as shown by synchrotron PCI. This result is in agreement with the expected location of zinc in the postsynaptic density (PSD) of hippocampal synapses where zinc could play a structural function in stabilizing proteins associated to the PSD such as Shank3 (Tao-Cheng et al., 2016).

Our data highlight the specific localization of copper within F-actin-rich regions along dendrites. The position, size and shape of the F-actin protrusions observed by STED microscopy (Fig. 2 and Fig S1–S3) are representative of dendritic spines (Chazeau & Giannone, 2016; Sala & Segal., 2014). Although actin-rich structures should predominantly be components of dendritic spines, other actin-rich structures may be present such as actin patches (Konietzny et al., 2017). Both dendritic spines and actin patches are important triggers of synaptic differentiation. It is noteworthy that copper distribution is only partially correlated with F-actin since copper is not detected in regions of lower SiR-actin fluorescence (Fig. 3). This observation could be explained by two different mechanisms. Copper might bind directly to F-actin but copper content is below the detection limit in regions containing lower amount of F-actin. Copper might not bind directly to F-actin but rather to other potential Cu-binding biomolecules that can interact with F-actin in specific areas of the dendritic protrusions where F-actin content is high. In an IMAC study focused on the copper proteome of HepG2 cells, 48 cytosolic proteins and 19 microsomal proteins displayed Cu-binding ability, among them γ-actin was identified both in the microsomal and the cytosolic fractions (Smith et al., 2004). Since γ-actin is also highly expressed in neurons, this IMAC result supports a direct binding of copper to F-actin in hippocampal neurons as well. In this same study, cofilin was also identified as putative Cu-binding protein in the cytosolic fraction (Smith et al., 2004). Copper could interact indirectly with molecules involved in the regulation of F-actin network in the dendritic spine such as cofilin, explaining the partial co-localization with F-actin. It is interesting to highlight that proteins binding copper with very high affinity (i.e. superoxide dismutase) are not separated by IMAC probably because the Cu-binding site is not vacant. Therefore IMAC results suggest a relatively labile Cu-binding to actin, or to cofilin, a mechanism in agreement with the potential function of Cu in the dynamic regulation of F-actin. It is also remarkable that α-tubulin and β-tubulin were identified as putative Cu-binding proteins in the IMAC study (Smith et al., 2004). The IMAC method may give positive results for metals of similar reactivity and does not necessarily reflect an interaction regulated by the cellular machinery. Because copper was not detected in dendritic shafts, our results suggest that if a physiologically relevant interaction existed between copper and tubulin it would be below the detection limit and therefore much less favorable than the one seen for zinc. The binding of α-tubulin and β-tubulin to copper in the IMAC experiments indicates the existence of a metal binding site, our study suggests that this metal could be Zn in living cells.

It is interesting to underline that iron was not detected within dendritic spines, meaning that iron content in this compartment is below the detection limit of the method, suggesting that contrary to copper and zinc, iron is probably not directly involved in the morphogenesis of the dendritic spines. On the other hand, iron was observed as local hot spots of few hundred of nanometers along dendrites, the exact nature of these local iron-enrichment remains to be elucidated. A possible explanation could be the presence of iron in mitochondria known to be present in dendrites of hippocampal neurons (Bastian et al., 2019).

From a methodological perspective, the combination of STED super resolution microscopy and nano-SXRF imaging opens numerous perspectives of application to investigate the chemical biology of metals. This correlative imaging stands as a solid new tool for the identification of metalloproteins directly in cells by correlating cellular imaging methods at a supramolecular scale. From a biological perspective, given the importance of cytoskeleton proteins in the morphological plasticity of neuronal connections (Sala & Segal., 2014), our results contribute to explain why copper and zinc are present in higher concentrations in the nervous system than in other organs. Our results indicate a broad role of copper and zinc in cytoskeleton architecture and provide a better understanding of metals functions in neurons, and may explain why metal dyshomeostasis, in particular copper or zinc depletion, are linked to numerous neuronal disorders.

## Methods

### Culture of primary rat hippocampal neurons

Primary rat hippocampal neurons were cultured on silicon nitride (SN) membranes (Silson Ltd) consisting in square silicon frames of 5 x 5 mm^2^ and 200 μm thickness with a central SN membrane of 1.5 x 1.5 mm2 and 500 nm thickness. During manufacturing, a second, smaller (0.1 x 0.1 mm^2^), SN membrane is added in one of the corners of the silicon frame to serve as orientation object. Primary rat hippocampal neurons were dissociated from E18 Sprague-Dawley rat embryos (Janvier labs) and plated on the SN membranes previously treated with 1 mg.ml^-1^ poly-lysine (Sigma) in a 0.1 M borate buffer pH 8.5. The SN membranes were placed on an astrocyte feeder layer growing on a dish treated by 0.1 mg.ml^-1^ poly-lysine in a 0.1 M borate buffer pH 8.5, in Neurobasal medium (Gibco) as described in the protocol from Kaech and Banker (2006). At 3-4 days *in vitro* (DIV3-4), cell cultures were treated with 2 μM of cytosine arabinofuranoside (Sigma) to limit the growth of the glial cells. From DIV6 and twice a week, half the Neurobasal medium was removed and replaced by BrainPhys medium (STEMCELL) at 300 mOsm, a culture medium designed to respect neuronal activity for *in vitro* models (Bardy et al., 2015). To develop dendritic spines, neurons were maintained in culture at 36.5°C in 5% CO_2_ atmosphere until DIV15.

### Cell labeling for STED and confocal imaging

For live-cell microscopy of tubulin and actin, fluorogenic probes based on silicone rhodamine (SiR) from Spirochrome were used according to manufacturer instructions and published protocols (Lukinavičius et al., 2014, Lukinavičius et al., 2016). For single color STED imaging, SiR-tubulin or SiR-actin were added to DIV15 neurons in the BrainPhys medium at a final concentration of 1 μM during 1.5 h at 37°C. For dual color imaging, SiR-tubulin and SiR700-actin were added to to the BrainPhys medium at 1 μM final concentration each and neurons were exposed during 1.5 h at 37°C.

### STED and confocal imaging

Confocal and STED microscopy were performed on a commercial Leica DMI6000 TCS SP8 X microscope. DIV15 neurons cultured on SN membranes and labeled with SiR fluorogenic probes were maintained in the microscope chamber at 37°C in a Tyrode’s solution (D-Glucose 1 mM, NaCl 135 mM, KCl 5 mM, MgCl_2_ 0.4 mM, CaCl_2_ 1.8 mM and HEPES 20 mM) pH 7.4 at 310 mOsm, the osmolarity of the BrainPhys culture medium. For live cell microscopy the SN membrane was mounted in a Ludin chamber. The SN membrane is placed on the glass coverslip of the Ludin chamber, with neurons facing the coverslip to minimize the distance between the objective and the cells to be observed. Confocal and STED images were acquired with a HC-PL-APO-CS2 93x immersion objective in glycerol with a numerical aperture of 1.3 and a scan speed of 400 Hz. SiR fluorogenic probes were excited at 640 nm (670 nm for SiR700) and the signal was detected with a Leica HyD hybrid detector with a window of emission recording from 651 to 680 nm for SiR and from 710 to 750 for SiR700. For STED acquisitions, the fluorescence outside the center was quenched with a 775 nm pulsed diode laser synchronized with excitation.

### STED spatial resolution

The confocal lateral resolution was calculated as 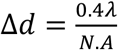 where Δ*d* is the smallest resolvable distance between two objects and N.A. is the numerical aperture. For confocal microscopy, the lateral resolution calculated was 194 nm. The STED lateral resolution was calculated as 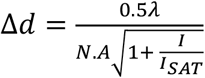 with λ the emission wavelength and 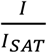 the saturation factor where *I* is the peak intensity of the depletion beam and *I_SAT_* the saturation intensity corresponding to the value of half emission signal. With a λ emission wavelength for SiR-tubulin of 674 nm, a saturation factor 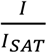 between 34.5 and 57.5, and a N.A. of 1.3, the smallest distance between two objects that could be resolved was included between 32 and 44 nm. To fully exploit the spatial resolution of the STED setup (between 32 and 44 nm), the pixel size of STED images was set to 25 nm for oversampling the data.

### Plunge-freezing and freeze-drying

Immediately after STED observations the neurons were plunge-frozen. Samples were quickly rinsed in a 310 mOsm ammonium acetate solution, pH 7.4 to remove extracellular inorganic elements present in Tyrode’s solution that would interfere with nano-SXRF element mapping. The osmolarity of Tyrode’s and ammonium acetate solutions were measured with a vapor pressure osmometer (VAPRO 5600, Elite) and adjusted to the initial values of the BrainPhys culture medium (310 mOsm). Then the cells were blotted with Whatman paper. To avoid primary specimen distortions, the blotting is performed without touching the cells, by capillarity from the back and from the side frame of the SN membranes. Samples are plunge-frozen during 20 seconds in 2-methylbutane (Sigma) cooled down at −165°C in liquid nitrogen. Excess 2-methylbutane was carefully blotted with Whatman paper cooled in liquid nitrogen vapors and samples transferred in the freeze-drier. Neurons were freeze-dried under gentle, minimally invasive conditions, during 2 days at −90°C and 0.040 mbar in a Christ Alpha 1-4 freeze drier, allowing preservation of cell structure as verified in our previous work (Perrin et al., 2015). Then the temperature and the pressure were slowly raised up to room temperature and ambient pressure and the samples were stored at room temperature within a desiccator until synchrotron analyzes.

### Synchrotron nano X-ray Fluorescence microscopy (SXRF) and phase contrast imaging

Synchrotron experiments were performed on the ID16A Nano-Imaging beamline at the European Synchrotron Radiation Facility (Grenoble, France) (Da Silva et al., 2017). The beamline is optimized for X-ray fluorescence imaging at 20 nm spatial resolution, as well as coherent hard X-ray imaging including in-line X-ray holography and X-ray ptychography. Featuring two pairs of multilayer coated Kirkpatrick-Baez (KB) focusing mirrors, the beamline provides a pink nanoprobe (Δ*E/E* ≈ 1%) at two discrete energies: *E* = 17 keV and 33.6 keV. Despite the larger focus at lower energy, for the correlative STED-SXRF experiment 17 keV was chosen as it is more efficient in exciting the X-ray fluorescence of the biologically relevant elements. The X-ray focus with dimensions of 35 nm (H) x 57 nm (V) provided a flux of 3.7 10^11^ ph/s. The focus spot size was determined with a lithographic sample consisting of a 10 nm thick, 20 x 20 μm2 square of nickel on a 500 nm thick SN membrane. The SN membranes holding the neurons were mounted in vacuum on a piezo nano-positioning stage with six short range actuators and regulated under the metrology of twelve capacitive sensors (Villar et al., 2018). An ultra-long working distance optical microscope was used to bring the sample to the focal plane (depth-of-focus ± 3 μm) and to position the STED regions of interest in the X-ray beam (see section sample positioning below). The samples were scanned with an isotropic pixel size of 40 nm, in some cases 20 nm for the scans of smaller size, and 100 ms of integration time. The X-ray fluorescence signal was detected with two custom energy dispersive detectors (Rayspec Ltd) at either side of the sample, holding a total of ten silicon drift diodes. The quantitative data treatment of the SXRF data was performed with Python scripts exploiting the PyMCA library (Solé et al., 2007), using the fundamental parameter method with the equivalent detector surface determined by calibration with a thin film reference sample (AXO DRESDEN GmbH). The resulting elemental areal mass density maps (units: ng.mm^-2^) were visualized with ImageJ. The X-ray phase contrast imaging (PCI) exploits in-line holography. It is performed on the same instrument moving the sample a few millimeters downstream of the focus and recording X-ray in-line holograms with a FReLoN CCD based detector located 1.2 m downstream of the focus (Mokso & Cloetens, 2007). X-ray holograms were collected in the Fresnel region at four different focus-to-sample distances to assure efficient phase retrieval. At each distance, images at 17 different lateral sample positions were recorded and averaged after registration, to eliminate artefacts related to the coherent mixing of the incident wavefront and the object. The neurons being pure and weak phase objects, phase retrieval was performed using the contrast transfer function approach (Cloetens et al., 1999), implemented in ESRF inhouse code using the GNU Octave language. The phase maps, proportional to the projection of the electron density, had a final pixel size of 15 nm and a field of view of 30 x 30 μm^2^. The spatial resolution was approximately 30 nm, similar to the STED and SXRF images.

### Sample positioning

During STED microscopy the orthonormal coordinates (x,y) of the regions of interest are recorded according to three reference positions on the SN frame, 3 corners of the square SN membrane. The first reference position corresponds to the corner close to the 0.1 x 0.1 mm^2^ orientation membrane, the second and third reference points to the corners selected from the first one in the clockwise direction. Using these 3 reference points the new coordinates (x’,y’) of the selected regions of interest can be calculated on the synchrotron ID16 setup using coordinate transformation equations.

### Calculation of SXRF LOD

The limit of detection (LOD) obtained with the ID16A nano-SXRF setup was determined for the elements phosphorus, sulfur, potassium and zinc according to IUPAC guidelines (IUPAC, 1997).

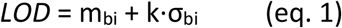

where m_bi_ is the mean of the blank measures, σ_bi_ is the standard deviation of the blank measures, and k is a numerical factor chosen according to the confidence level desired, k=3 for LOD. The resulting LOD values for each element based on the mean and standard deviation of 12 different blank analyses are presented in Table 1.

### Zinc chelation with TPEN

TPEN (Sigma Aldrich) cytotoxicity was controlled using the ReadyProbes cell viability imaging kit, Blue/Green, from Thermo-Fischer Scientific (R37609), according to the manufacturer instructions. Mature neurons (15 DIV) were treated with increasing concentrations of TPEN (0 to 10 μM), dissolved in analytical grade ethanol (Sigma Aldrich) to avoid trace element contamination, at a final ethanol concentration of 0.1% (v/v) in the culture medium, during 24h. Zinc and other chemical elements content was monitored in exposure solutions by PIXE (Particle Induced X-ray Emission) analysis. The counting of total and dead neurons was performed using fluorescence imaging on a Leica DM5000 microscope. The concentration of 5 μM TPEN was selected for further zinc chelation experiments since this concentration induced a subcytotoxic effect, with 83% of neurons alive compared to the control group (100%). Three experimental groups were compared by quantitative fluorescence microsocpy, neurons exposed to 5 μM TPEN for 24h, neurons exposed simultaneously to 5 μM TPEN and 10 μM ZnCl_2_ for 24h, and control neurons exposed to the vehicle solution of 0.1% (v/v) ethanol in culture medium.

### Fluorescence labeling of tubulin and F-actin

Primary rat hippocampal neurons were cultured on glass coverslip on an astrocyte layer following the method developed by Kaech & Banker (2006). Mature neurons (15 days *in vitro*) were exposed to 5 μM of TPEN during 24 h. Primary neurons were fixed 20 minutes at room temperature in 4% paraformaldehyde, 0.25% glutaraldehyde and 0.1% Triton-X100 in a cytoskeleton buffer with a final concentration of 10 mM MES (2-(N-morpholino)ethanesulfonic acid), 150 mM NaCl, 5 mM EGTA (ethylene glycol-bis(β-aminoethyl ether)-N,N,N’,N’-tetraacetic acid), 5 mM glucose and 5 mM MgCl_2_ (pH 6,2). After rinsing in the same cytoskeleton buffer then in PBS (phosphate-buffered saline) the aldehyde fluorescence was quenched in 0.1 % NaBH4 in PBS during 5 minutes. After 5 minutes in 0.2% Triton-X100 the neurons were rinced in PBS and then blocked in 2% BSA in PBS during 45 min. Neurons were incubated 45 min with the primary antibody Mouse IgG1 α beta tubulin (Sigma Aldrich, T4026) diluted in 2% BSA (1:3000 v/v) and blocked again 15 min in the 2% BSA solution. Neurons were incubated 30 minutes in darkness with phalloïdin-Alexa647 (ThermoFisher Scientific, A22287) (1:40; v/v) and Goat anti Mouse IgG1 CF568 (Ozyme, BTM20248) (1:500, v/v). Coverslips were rinced in PBS and neurons were fixed again in 2% paraformaldehyde in PBS for 10 minutes to stabilize the phalloïdin before to be treated with 50 mM NH4Cl to quench aldehyde fluorescence.

### Quantitative fluorescence microscopy and data analysis

Microscopy images of F-actin and β-tubulin of neurons were collected the day after chemical fixation with a Leica DM5000 upright fluorescence microscope at 63x magnification. All fluorescence images were acquired on the same day using the same experimental conditions. For each experimental group (control, 5 μM TPEN and 5 μM TPEN + 10 μM Zn), between 2 and 4 coverslips were analyzed. For each coverslip an average of 20 images were recorded randomly. The experiments were repeated three times on neurons dissected from three different animals to ensure biological reproducibility. Data were treated using ImageJ software (http://imagej.nih.gov/ij/). F-actin and β-tubulin segmentation of the images was performed using histogram thresholding. The threshold value was defined as the value immediately greater than the bin value with the maximum counts in the histogram. Neuronal somas were removed from the images using clear selection to select only dendrite areas. Mean fluorescence intensities (counts/pixel) were calculated for each channel (F-actin and β-tubulin) on each segmented image. Mean fluorescence intensities values per image of 70 μm x 100 μm were then used to plot the Factin and β-tubulin intensity signal for each of the three experimentals groups (control, 5 μM TPEN and 5 μM TPEN + 10 μM Zn). Fluorescence intensity was normalized against the median intensity of the corresponding control group as presented in Fig. 6 and S6.

### Statistical analysis

For nano-SXRF data analysis of element distrbutions, Pearson correlation coefficients were calculated using R software (R Core Team, 2019) with rcmdr package (Fox & Bouchet-Valat, 2019). For fluorescence microscopy experiments, fluorescence intensity was normalized against the median of the control group for each experiment. Normality of distribution and homogeneity of variances were respectively checked with Shapiro-Wilk test and Bartlett’s test. Comparison of normalized intensity between two independent groups was performed using t-test or Mann-Whitney test if normal distribution or homogeneity of variance were not met. ANOVA was used for comparison of three groups, with Tukey post-hoc test for pairwise comparisons. Kruskal-Wallis test was run when assumptions of normality or homogeneity of variances were not met. Significant Kruskal-Wallis test (p<0.01) was followed by Dunn’s test with p-values adjusted for pairwise comparisons (Holm’s method). Statistical analysis was made using Rstudio v 1.2.5001 (RStudio Team, 2015), R software v3.6.1 (R Core Team, 2019) with rcmdr (Fox & Bouchet-Valat, 2019), tidyverse (Wickham et al., 2019), ggpubr (https://rpkgs.datanovia.com/ggpubr/), rstatix (https://rpkgs.datanovia.com/rstatix/) packages. For details see Table S2.

## Acknowledgments

This project was supported by a doctoral fellowship from the University of Bordeaux (F.D.), a grant from Centre National de la Recherche Scientifique (CNRS) through the MITI interdisciplinary program (R.O.), a PEPS grant from CNRS and IDEX Bordeaux (R.O. and D.C.), ERC grant ADOS (339541) and DynSynMem (787340) to D.C. and support from the Regional Council Nouvelle Aquitaine. SXRF experiments were peformed at the European Synchrotron Radiation Facility (ESRF), Grenoble, France. We are grateful to ESRF staff for assistance in using ID16A beamline. STXM experiments were performed at SOLEIL synchrotron, Gif-sur-Yvette, France. We acknowledge SOLEIL for provision of synchrotron radiation facilities and we would like to thank HERMES staff and F. Porcaro from CENBG, for their help during the experiments. We acknowledge P. Mascalchi and C. Poujol from Bordeaux Imaging Center, part of the France BioImaging national infrastructure, for support in microscopy; C. Breillat and N. Retailleau from the Cell Biology facility of IINS for neuronal cell culture.

## Author contributions

F.D., D.C., and R.O. designed the experiments. F.D. and E.V. designed and performed protocols for cell cultures. F.D., S.R., P.C., and R.O. performed synchrotron radiation experiments. P.C. developed nano-SXRF and PCI methodology. F.D., A.C., and P.C. performed data treatment of synchrotron radiation experiments. F.D. and R.O. performed STED experiments. F.D. analyzed STED experiments and S.R. provided advice. F.D. performed fluorescence microscopy experiments. S.R. performed statistical analysis. F.D., A.C., D.C. and R.O. wrote the manuscript.

## Data availability

The data supporting the findings of this study are available from the corresponding author upon request. Synchrotron data are managed according to ESRF policy based on the PaNdata Data Policy.

## SUPPLEMENTARY MATERIALS

**Fig. S1.**
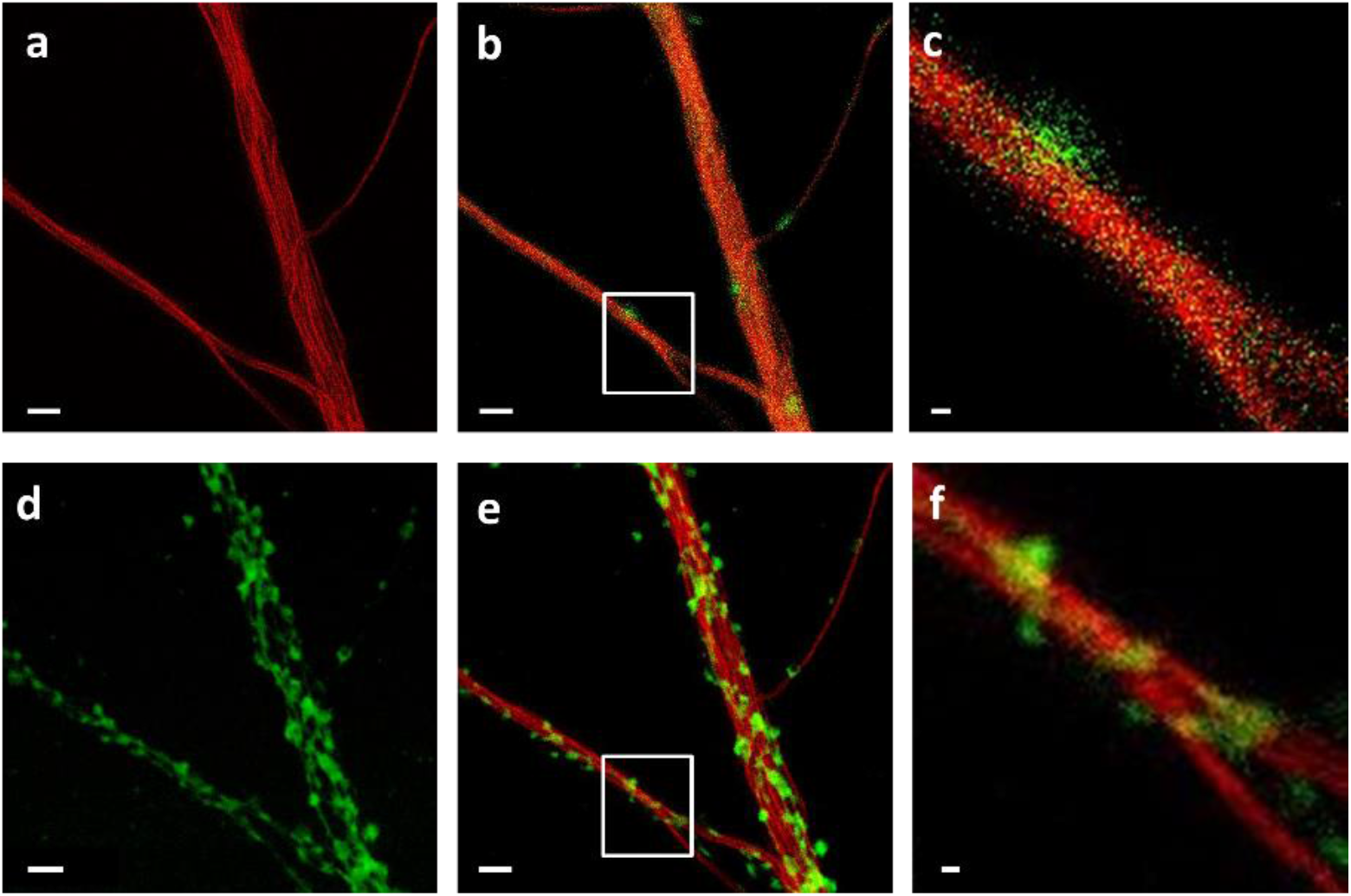
Correlation of STED imaging on live cells and of STXM microscopy after cryofixation and freeze-drying. a) Live STED imaging of dendrites after SiR-tubulin labeling. b) Confocal image of the same dendrites showing tubulin in red and F-actin in green (SiR700-actin labeling). c) Zoom on the framed region of image b) showing a F-actin-rich protrusion (in green) on the tubulin-rich dendrite (in red). d) STXM image of the same dendrite obtained with a spatial resolution of 25 nm after cryofixation and FD. e) Superposition of live cell STED (red) and FD cell STXM (green) images showing the good preservation of the cell structures after cryofixation and FD. F) Zoom on the framed region in the image e) showing the presence of dendritic protrusions revealed by STXM (in green). One of these protrusions is co-localized with the F-actin fluorescence visible in c). Scale bars = 2 μm except for images c and f, scale bars = 250 nm.

**Fig. S2.**
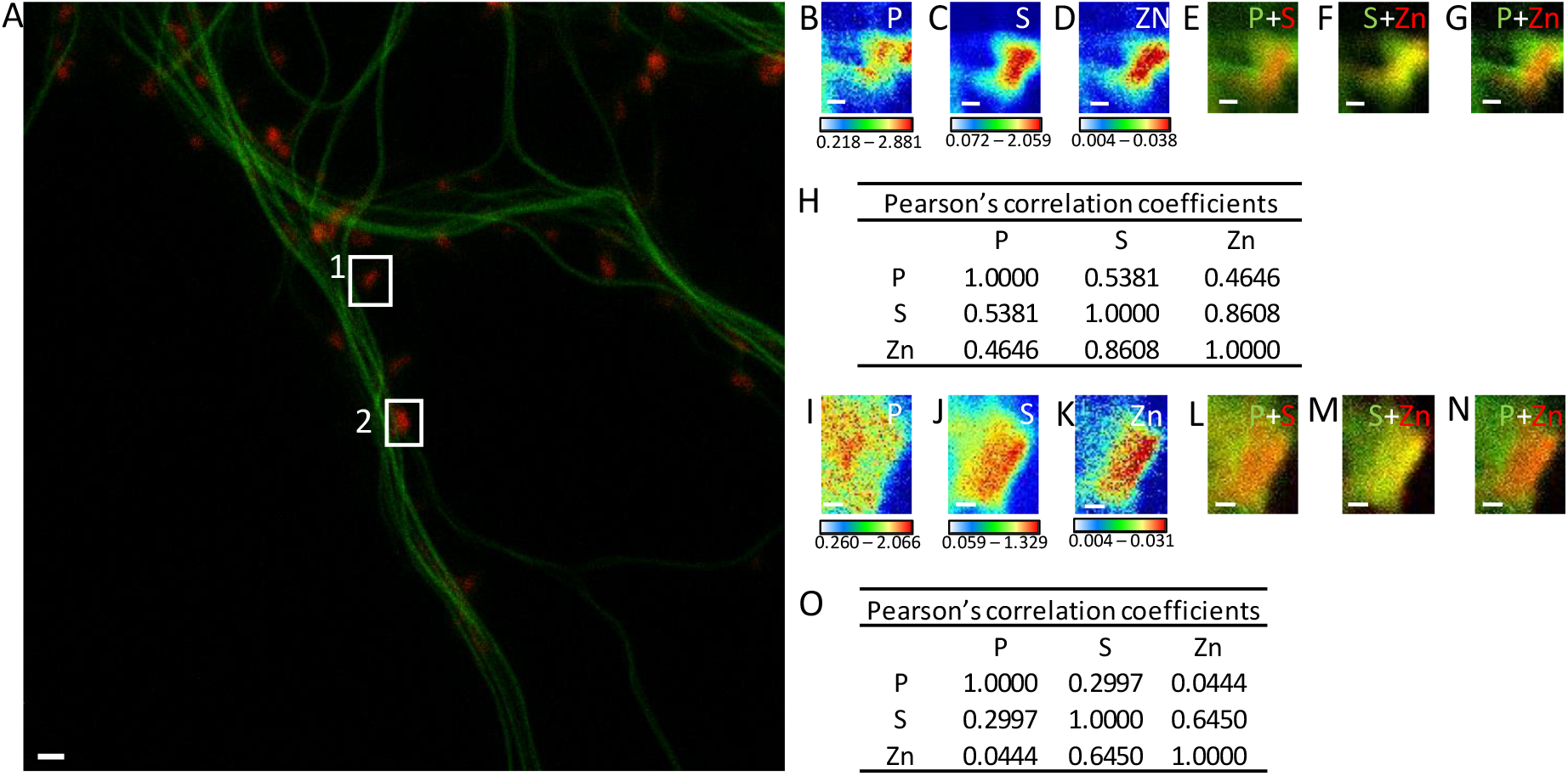
Element distributions in dendritic spines. (A) STED image of SiR-tubulin (green) and confocal image of SiR700-actin (red). (B-D) SXRF element maps (P, S, Zn) from spine framed in (A) as region 1. (E) Overlay image of phosphorus (green) and sulfur (red). (F) Overlay image of sulfur (green) and zinc (red). (G) Overlay image of phosphorus (green) and zinc (red). (H) Pearson’s correlation coefficients for the elements in spine 1. (I-K) SXRF element maps (P, S, Zn) from spine framed in (A) as region 2. (L) Overlay image of phosphorus (green) and sulfur (red). (M) Overlay image of sulfur (green) and zinc (red). (N) Overlay image of phosphorus (green) and zinc (red). (O) Pearson’s correlation coefficients for the elements in spine 2. Scale bars 200 nm except for (A) 1 μm. Color scales: min-max values in ng.mm^-2^.

**Fig. S3.**
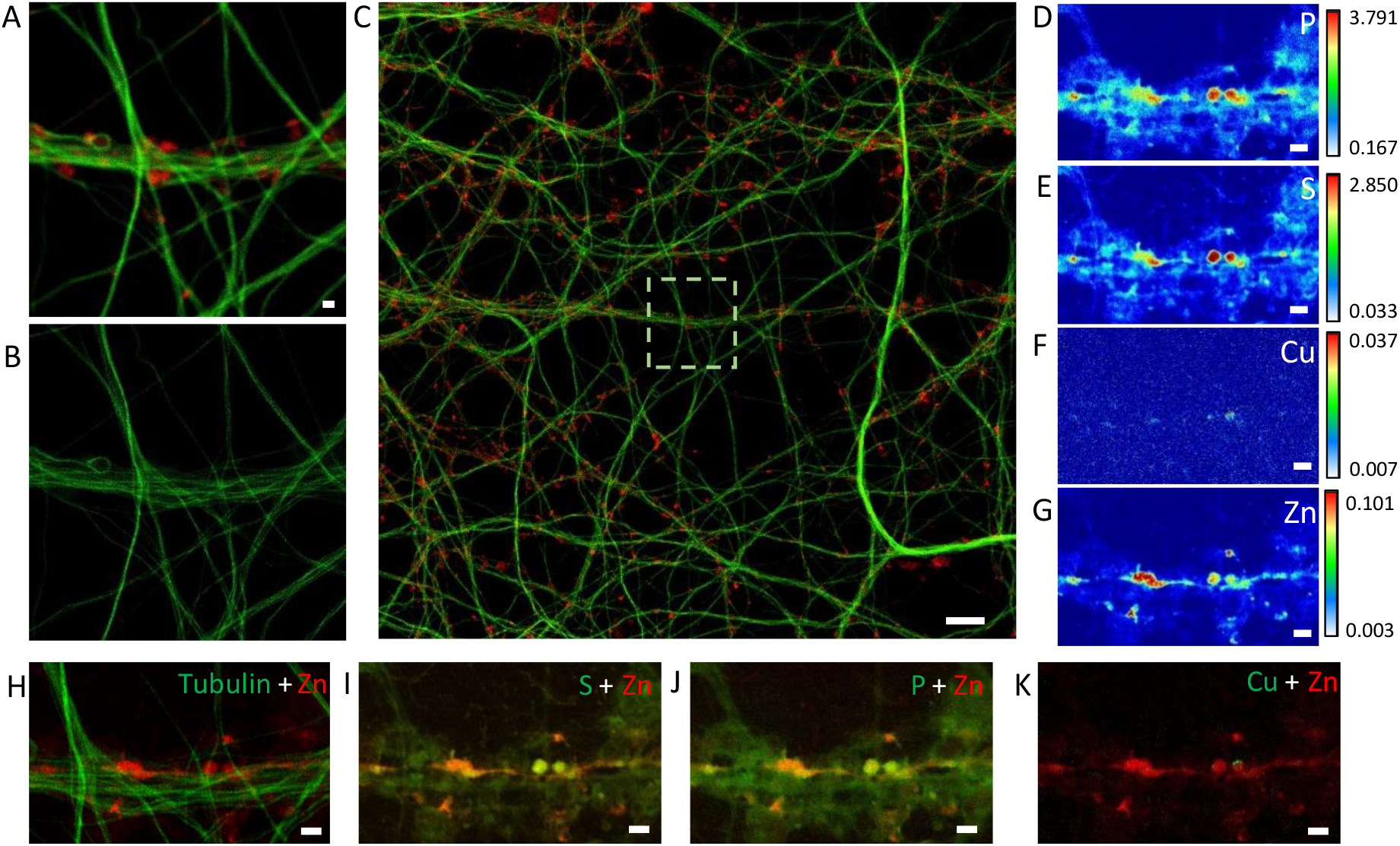
Element distributions in dendrites and spines. (A) Confocal image of SiR-tubulin (green) and SiR700-actin (red) in dendrites. (B) STED image of SiR-tubulin in the same zone as in (A). (C) Confocal large image including the region shown in (A) with SiR-tubulin (green) and SiR700-actin (red). (D-G) SXRF element maps (P, S, Cu, Zn) from the same region as (A). (H) Overlay image of STED SiR-tubulin (green) and zinc (red). (I) Overlay image of sulfur (green) and zinc (red). (J) Overlay image of phosphorus (green) and zinc (red). (K) Overlay image of copper (green) and zinc (red). Scale bar: 1 μm except for (C) 10 μm. Color scales: min-max values in ng.mm^-2^.

**Fig. S4.**
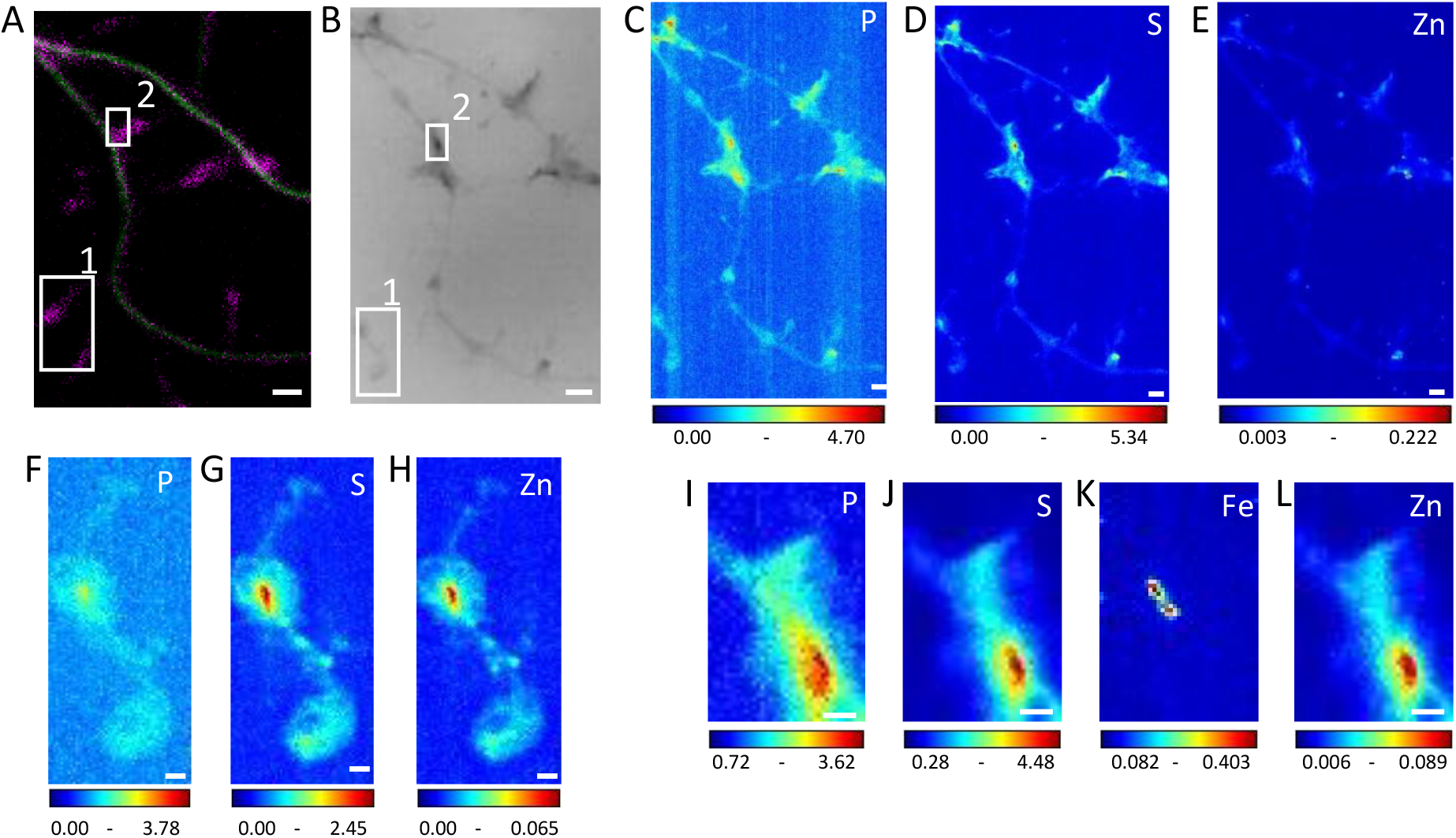
Multimodal imaging of dendrites and spines. (A) STED SiR-tubulin (green) and confocal SiR700-actin (magenta) (B) Synchrotron radiation X-ray PCI of the same area as (A). (C-E) SXRF element maps (P, S, Zn) from the same region as (A) and (B). (F-H) SXRF element maps (P, S, Zn) from the framed region 1 in (B). (I-L) SXRF element maps (P, S, Fe, Zn) from the framed region 2 in (B). Scale bars: 200 nm, except (A) and (B) 1 μm. Color scales min-max values in ng.mm^-2^.

**Fig. S5.**
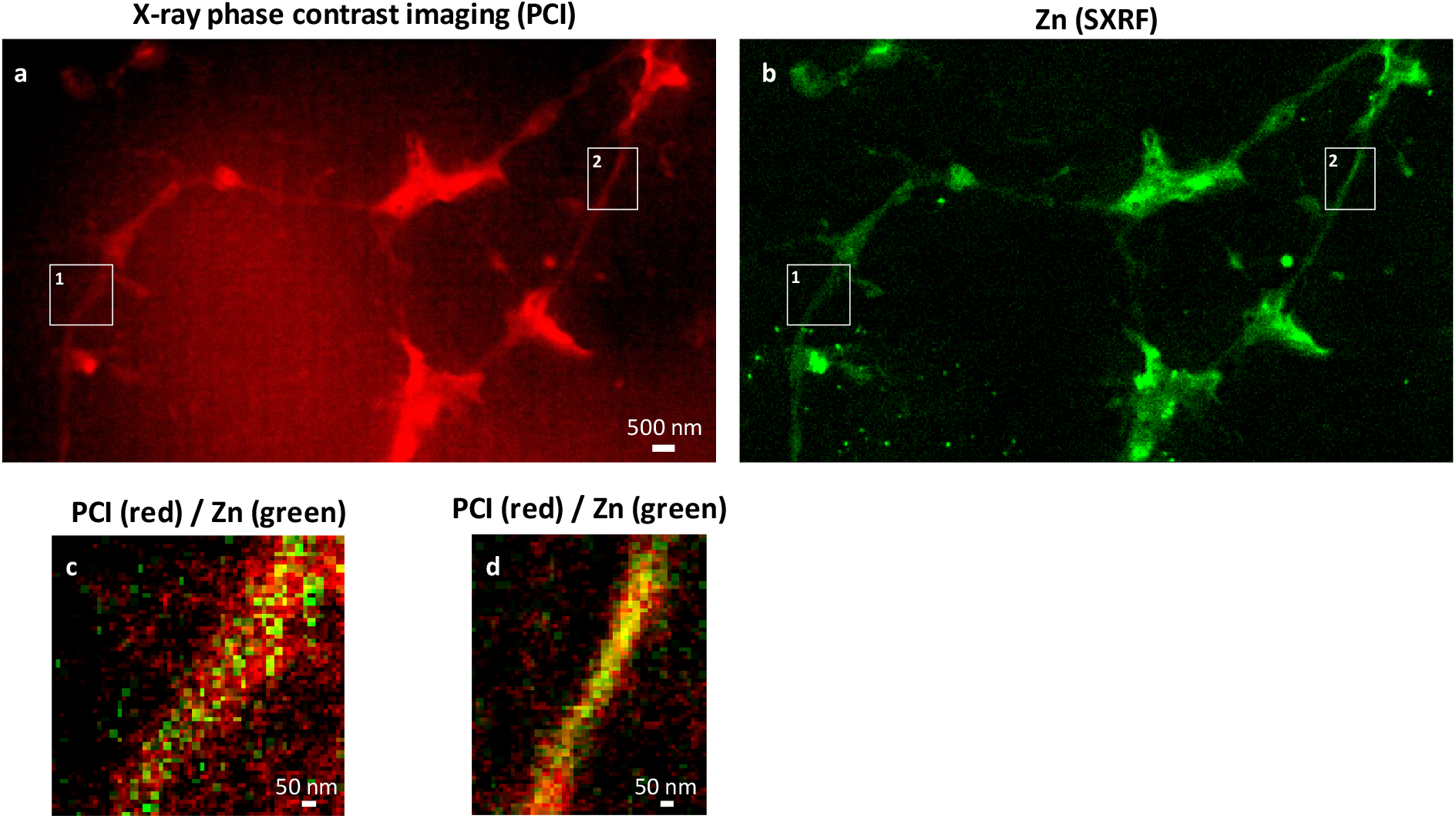
a) Synchrotron X-ray phase contrast imaging (PCI) showing the dendrites and spines morphology. b) SXRF imaging of Zn in the same sample area. c) Overlay image of PCI (red) and Zn (green) from a ‘thick’ dendrite (region 1) showing Zn inside the dendritic structure. d) Overlay image of PCI (red) and Zn (green) from a ‘thin’ dendrite (region 2) showing that Zn distribution and dendrite morphology have the same dimensions.

**Fig. S6.**
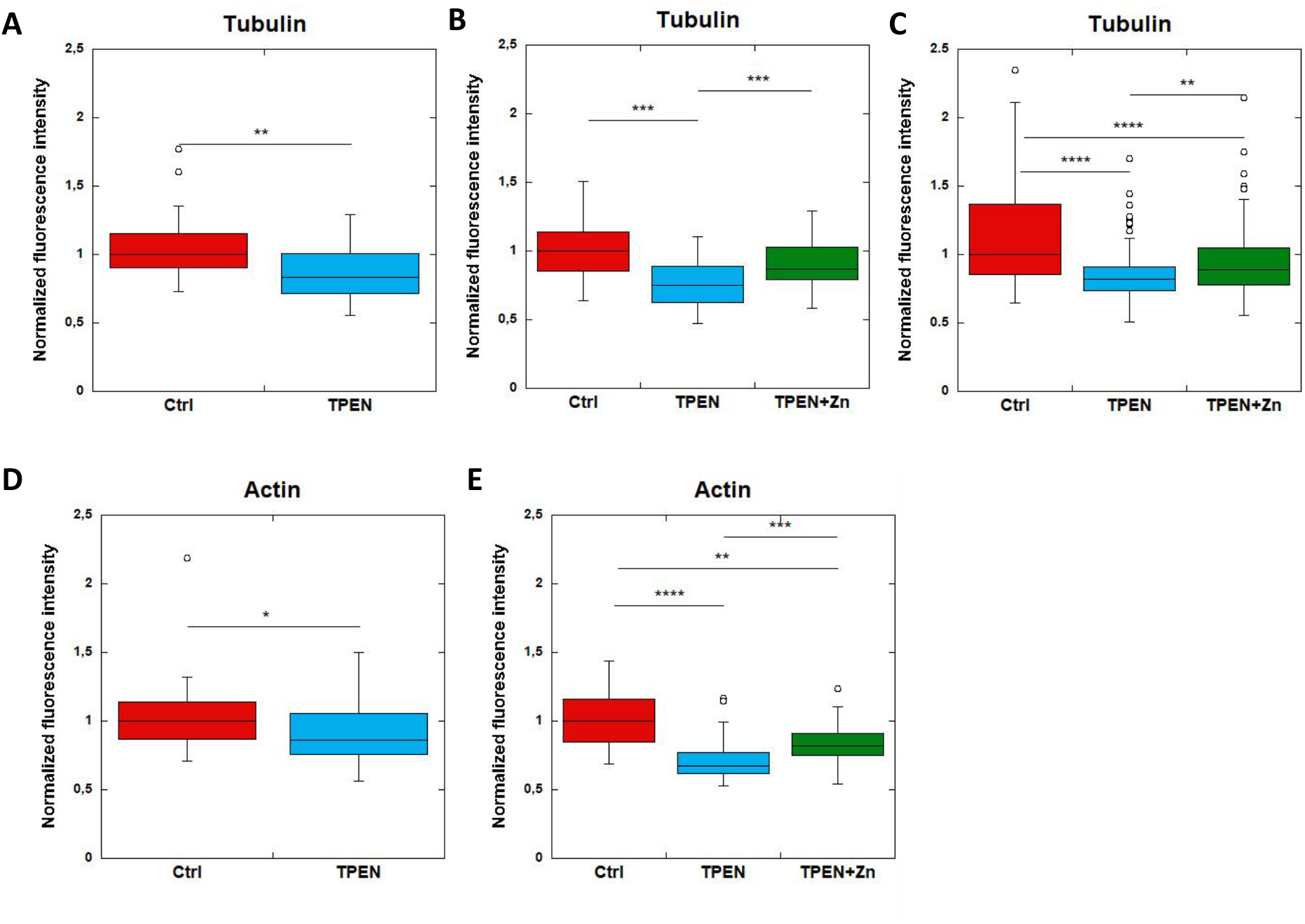
F-actin and β-tubulin fluorescence in control, TPEN, or TPEN and Zn treated neurons. Replicates of TPEN (5 μM, 24h) or TPEN (5 μM, 24h) and Zn (10 μM, 24h) exposure experiments presented in fig. 6. (A-C) Comparison of β-tubulin normalized fluorescence intensities. (D-E) Comparison of F-actin normalized fluorescence intensities. *p-value <0.05, **p-value <0.001, ***p-value <0.001, ****p-value <0.0001 (see Table S2 for details on statistical analysis).

**Table S1.** Nano-SXRF quantitative data. Analysis of chemical elements content for 21 regions showing zinc and tubulin co-localization, expressed in ng.mm^-2^ and in atoms.nm^-2^ (mean ± SD, n=21).

|  | Phosphorus | Sulfur | Potassium | Zinc |
| --- | --- | --- | --- | --- |
| ng.mm<sup>-2</sup> | 0.600 ± 0.085 | 0.211 ± 0.068 | 0.153 ± 0.034 | 0.0105 ± 0.005 |
| atoms.nm<sup>-2</sup> | 11.81 ± 3.29 | 3.96 ± 1.28 | 2.57 ± 1.23 | 0.09 ± 0.01 |

**Table S2.**
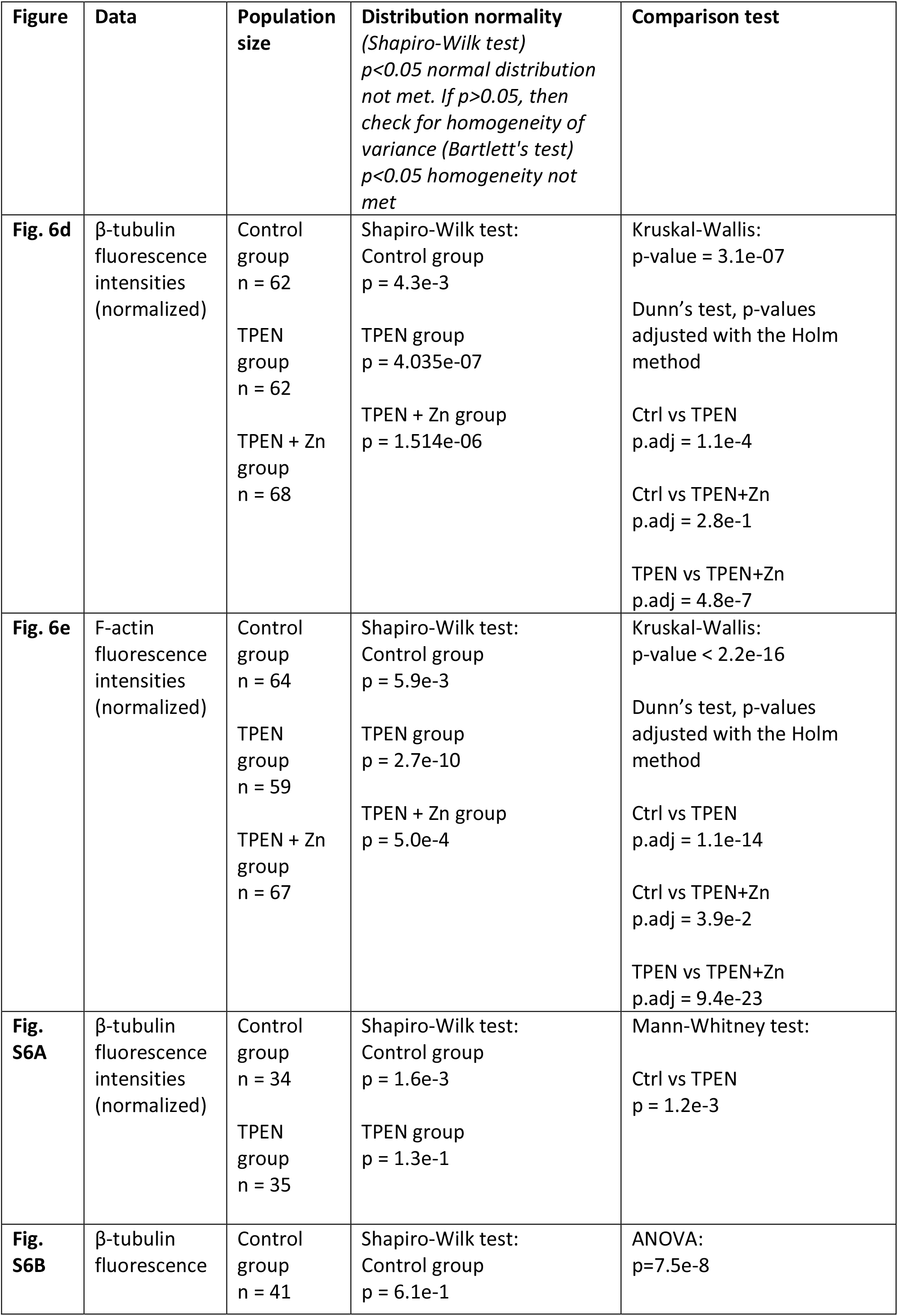
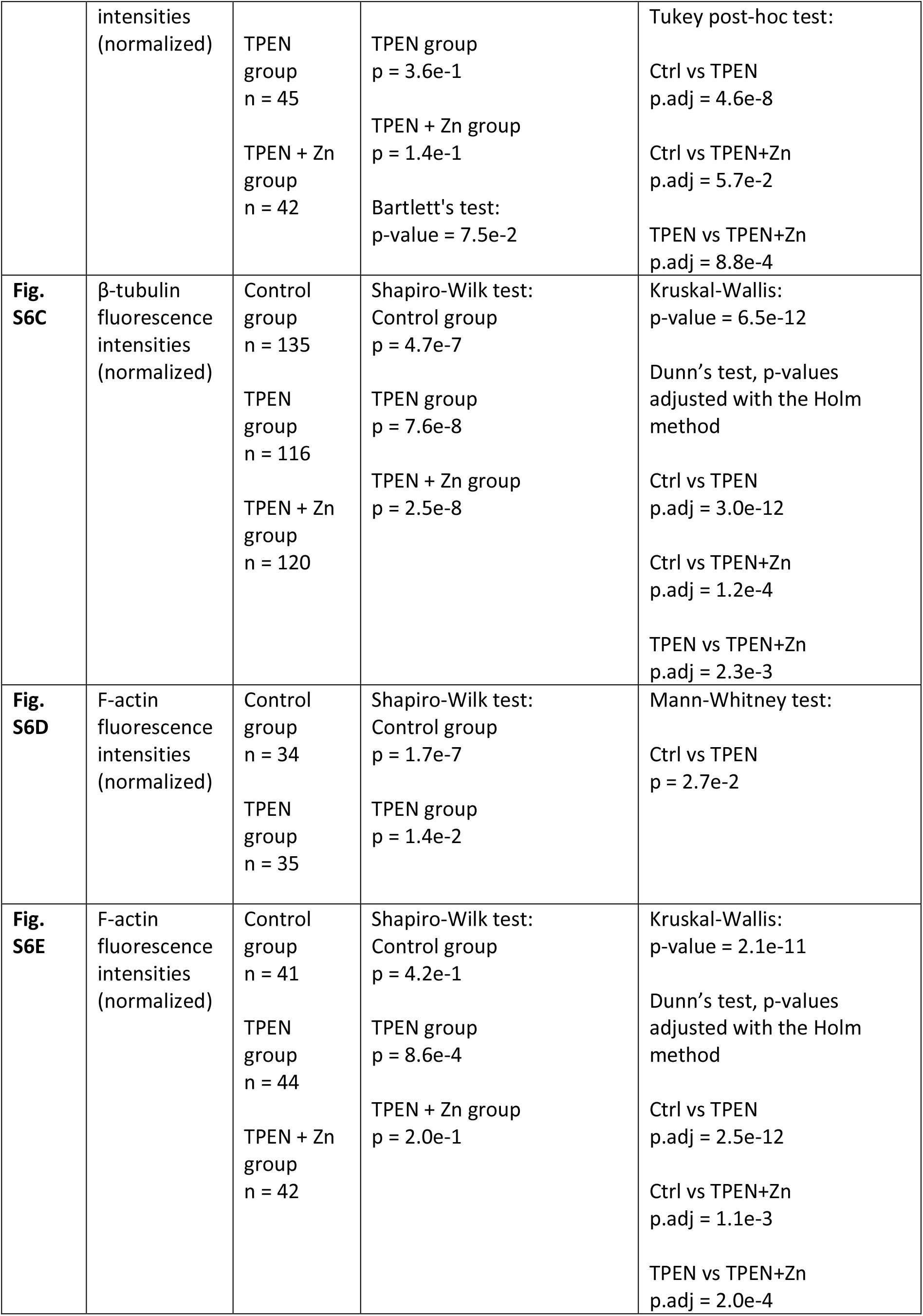
Statistical analysis of data.

| Figure | Data | Population size | Distribution normality<br>(Shapiro-Wilk test)<br>*p*<0.05 normal distribution not met. If *p*>0.05, then check for homogeneity of variance (Bartlett's test)<br>*p*<0.05 homogeneity not met | Comparison test |
| --- | --- | --- | --- | --- |
| **Fig. 6d** | β-tubulin fluorescence intensities (normalized) | Control group<br>n = 62<br>TPEN group<br>n = 62<br>TPEN + Zn group<br>n = 68 | Shapiro-Wilk test:<br>Control group<br>*p* = 4.3e-3<br>TPEN group<br>*p* = 4.035e-07<br>TPEN + Zn group<br>*p* = 1.514e-06 | Kruskal-Wallis:<br>*p*-value = 3.1e-07<br>Dunn's test, *p*-values adjusted with the Holm method<br>Ctrl vs TPEN<br>*p*.adj = 1.1e-4<br>Ctrl vs TPEN+Zn<br>*p*.adj = 2.8e-1<br>TPEN vs TPEN+Zn<br>*p*.adj = 4.8e-7 |
| **Fig. 6e** | F-actin fluorescence intensities (normalized) | Control group<br>n = 64<br>TPEN group<br>n = 59<br>TPEN + Zn group<br>n = 67 | Shapiro-Wilk test:<br>Control group<br>*p* = 5.9e-3<br>TPEN group<br>*p* = 2.7e-10<br>TPEN + Zn group<br>*p* = 5.0e-4 | Kruskal-Wallis:<br>*p*-value < 2.2e-16<br>Dunn's test, *p*-values adjusted with the Holm method<br>Ctrl vs TPEN<br>*p*.adj = 1.1e-14<br>Ctrl vs TPEN+Zn<br>*p*.adj = 3.9e-2<br>TPEN vs TPEN+Zn<br>*p*.adj = 9.4e-23 |
| **Fig. S6A** | β-tubulin fluorescence intensities (normalized) | Control group<br>n = 34<br>TPEN group<br>n = 35 | Shapiro-Wilk test:<br>Control group<br>*p* = 1.6e-3<br>TPEN group<br>*p* = 1.3e-1 | Mann-Whitney test:<br>Ctrl vs TPEN<br>*p* = 1.2e-3 |
| **Fig. S6B** | β-tubulin fluorescence | Control group<br>n = 41 | Shapiro-Wilk test:<br>Control group<br>*p* = 6.1e-1 | ANOVA:<br>*p*=7.5e-8 |
|  | intensities<br>(normalized) | TPEN<br>group<br>n = 45<br>TPEN + Zn<br>group<br>n = 42 | TPEN group<br>p = 3.6e-1<br>TPEN + Zn group<br>p = 1.4e-1<br>Bartlett's test:<br>p-value = 7.5e-2 | Tukey post-hoc test:<br>Ctrl vs TPEN<br>p.adj = 4.6e-8<br>Ctrl vs TPEN+Zn<br>p.adj = 5.7e-2<br>TPEN vs TPEN+Zn<br>p.adj = 8.8e-4 |
| **Fig. S6C** | β-tubulin<br>fluorescence<br>intensities<br>(normalized) | Control<br>group<br>n = 135<br>TPEN<br>group<br>n = 116<br>TPEN + Zn<br>group<br>n = 120 | Shapiro-Wilk test:<br>Control group<br>p = 4.7e-7<br>TPEN group<br>p = 7.6e-8<br>TPEN + Zn group<br>p = 2.5e-8 | Kruskal-Wallis:<br>p-value = 6.5e-12<br>Dunn's test, p-values<br>adjusted with the Holm<br>method<br>Ctrl vs TPEN<br>p.adj = 3.0e-12<br>Ctrl vs TPEN+Zn<br>p.adj = 1.2e-4<br>TPEN vs TPEN+Zn<br>p.adj = 2.3e-3 |
| **Fig. S6D** | F-actin<br>fluorescence<br>intensities<br>(normalized) | Control<br>group<br>n = 34<br>TPEN<br>group<br>n = 35 | Shapiro-Wilk test:<br>Control group<br>p = 1.7e-7<br>TPEN group<br>p = 1.4e-2 | Mann-Whitney test:<br>Ctrl vs TPEN<br>p = 2.7e-2 |
| **Fig. S6E** | F-actin<br>fluorescence<br>intensities<br>(normalized) | Control<br>group<br>n = 41<br>TPEN<br>group<br>n = 44<br>TPEN + Zn<br>group<br>n = 42 | Shapiro-Wilk test:<br>Control group<br>p = 4.2e-1<br>TPEN group<br>p = 8.6e-4<br>TPEN + Zn group<br>p = 2.0e-1 | Kruskal-Wallis:<br>p-value = 2.1e-11<br>Dunn's test, p-values<br>adjusted with the Holm<br>method<br>Ctrl vs TPEN<br>p.adj = 2.5e-12<br>Ctrl vs TPEN+Zn<br>p.adj = 1.1e-3<br>TPEN vs TPEN+Zn<br>p.adj = 2.0e-4 |

